# Myeloid cell IL-15 production in the brain supports Bystander CD8+ T-Cell Neuropathic Immune Responses following Virus infection

**DOI:** 10.1101/2025.05.13.653736

**Authors:** Jennifer N. Berger, Alayna Rosales, Dustin Heiden, Brendan Monogue, J. David Beckham, Leslie Berg

## Abstract

The central nervous system (CNS) includes a uniquely regulated immune response that supports homeostasis, response to injury, and response to pathogens. Recent work has shown that virus-associated immune responses in the CNS may contribute to neuronal injury and long-term outcomes such as neurocognitive decline. However, the fundamental mechanisms that regulate acute infiltration of immune cells from vascular compartments into the CNS are not well defined. Using an attenuated Venezuelan equine encephalitis virus TC83 (referred to as TC83) to inoculate using olfactory and intracranial injections, we show that infection in the CNS and olfactory pathways results in rapid infiltration of both CD4+ and CD8+ T-cells as early as 3- and 5-days post-infection. CNS-infiltrating CD8+ T-cells exhibit a bystander, memory phenotype (CD49d+, Tbet+, NKG2D+, Eomes+), are cytotoxic, and are recruited independent of antigen specific responses. We show that infiltration of CD8+ bystander T-cells is supported by microglia and infiltrating macrophage expression of IL-15 and interferon expression in the CNS. These innate antiviral immune signals support activation of bystander CD8+ T-cells in the CNS that contribute to tissue injury independent of virus replication at early time points post-infection. These data support a mechanism by IL-15 stimulates bystander memory CD8+ T-cells to enter the CNS and contribute to injury independent of antigen-specific stimulation.

**Importance:** Prior studies have shown that virus infections in the respiratory and olfactory nerve pathways can result in long term injury in the brain. However, the mechanisms that link virus infection in the olfactory neurons and brain injury are not understood. We show that virus infection of olfactory neurons results in immune stimulation in the brain of resident immune cells to release a cytokine called IL-15 and interferon. This results in infiltration and activation of non-specific T-cells that cause injury of neurons in the brain. This may be an important mechanism by which respiratory viruses and other viruses cause inflammation and injury in the brain.

## Introduction

The central nervous system (CNS) is a unique site of immune regulation largely defined by the blood-CNS barrier, the blood-cerebral spinal fluid barrier, and the pial-CNS barrier.(1) These barriers provide homeostatic regulation of CNS transport of small molecules while restricting CNS entry of large molecules, cells, and infectious agents such as viruses. The nervous system was traditionally thought to act independently of an organism’s immune response and be an “immune privileged site”. Increasing evidence has shown that the nervous system is not immune privileged but instead has a unique immune response that is critical for maintaining homeostasis and responding to pathogens.(2)

Recent studies have increasingly reported associations between infectious events and cognitive decline.(3–6) In AD cases, the discovery of herpes simplex virus (HSV) DNA in postmortem AD brains initiated a field of study to determine potential viral causes of AD, and several studies have linked virus gene expression in the brain with risk of developing AD.(4, 7–10) With the large COVID-19 pandemic, recent data also show that respiratory viruses like SARS-CoV2 can be associated with long-term cognitive decline and exacerbate dementia in patients with underlying neurodegenerative diseases.(11, 12) Similarly, there are increasing data linking infections and risk for PD through unclear mechanisms.(13–15) Recent data have shown that alpha-synuclein expression is critical in protecting neurons from viral infection in the CNS by modulating type 1 interferon signaling in the brain.(16, 17)

Viral infections in the upper airway are known to cause changes in neuroimmune responses in olfactory neurons and can cause direct infection of olfactory pathways resulting in modulation of innate immune responses in the olfactory bulb of the brain.(18–21) Olfactory changes and subsequent neurocognitive changes can be associated with acute viral infections in the olfactory pathway. Multiple studies have also shown that olfactory pathway impairments occur in approximately 90% of patients with Alzheimer’s Disease (AD) and Parkinson’s disease (PD), and olfactory impairment is associated with cognitive decline.(22–24) Additional studies have shown that olfactory deficits including impaired odor detection and odor discrimination manifest early in the course of neurodegeneration, preceding cognitive and motor symptoms by many years.(22, 25) In a two year longitudinal study of adults >65 years of age, cognitive decline was greater for individuals with olfactory dysfunction and carried at least one apolipoproteinEe4 allele compared to controls.(26) Olfactory bulb tissue from patients with PD exhibit increased gene expression of cytokine signaling pathways and MAPK signaling pathways compared to control tissue suggesting that neuroimmune responses may play a role in olfactory dysfunction related to neurodegenerative diseases.(27) While virus-induced changes in innate immune responses in olfactory pathways may overlap with the pathogenesis of neurodegeneration, the potential shared or overlapping mechanisms of injury are not known.

Taken together, these studies suggest that antiviral neuroimmune mechanisms may play a critical role in long term risk of neurodegenerative diseases. However, the fundamental mechanisms that regulate acute infiltration of immune cells from vascular compartments into the CNS are not well defined, and further studies defining these mechanisms may provide unique insights into the pathogenesis of neurodegeneration.

The cellular components of the neurovascular unit include the endothelium, pericytes, perivascular macrophages, and microglia (the CNS-resident macrophages).(28) Pericytes regulate leukocyte trafficking across endothelial cells and regulate T-cells through production of transforming growth factor-beta (TGFb) and retinoic acid. T-cell entry at the blood-CNS barrier is regulated at the glia limitans resulting in the classical perivascular ‘cuff’ pattern noted on pathology samples of the brain during acute inflammation.(28) CSF memory T-cells display expansion of restricted clones in the absence of neuroinflammation, and a brain-resident T-cell population consists of memory phenotypes predominantly from CD4+ and CD8+ memory T-cell subsets.(29, 30) Human white matter-derived resident brain CD8+ T-cells are characterized by low expression of transcription factors T-bet and eomesodermin (eomes).(31) The mechanism of memory T-cell trafficking to the CNS during acute viral infection, the role of T-cell receptor-dependent activation, and the mechanism of T-cell memory trafficking into the CNS during acute viral infection are not known.

Here, we sought to investigate the acute immune response in the CNS following infection with an attenuated vaccine strain of Venezuelan equine encephalitis virus TC83 (referred to as TC83). TC83 replicates to high titers in the brains of wild-type mice without lethality, thus enabling us to investigate the innate cellular immune response to viral infection. Using TC83, we show that infection in the CNS and olfactory pathways results in rapid infiltration of both CD4+ and CD8+ T-cells as early as 3- and 5-days post-infection. Moreover, the CNS-infiltrating CD8+ T-cells exhibit a bystander, memory phenotype (CD49d+, Tbet+, NKG2D+, Eomes+), are cytotoxic, and are recruited independent of antigen specific responses. We also show that infiltration of CD8+ bystander T-cells is supported by microglia and infiltrating macrophage expression of IL-15 as well as type I interferon. Following IL-15 and interferon-dependent entry to the CNS, bystander CD8+ T-cells contribute to tissue injury independent of virus replication.

## Results

### VEEV-TC83 Infection Induces Acute Infiltration of T-cells in the Olfactory bulb

We first determined the role of acute virus infection with TC83 in the recruitment of lymphocytes to the brain at early time points post-infection and prior to antigen-specific responses. Wild-type B6/C57 mice were injected with mock inoculum or VEEV-TC83 (10^5^pfu) by intracranial (i.c) inoculation. Brain tissue was harvested at day 5 post-infection and prepared for spectral flow cytometry to analyze lymphocyte subsets (**Fig. S1**). We found that VEEV-TC83 infection induced a significant increase in infiltration of CD4+ and CD8+ T-cells at day 5 post-infection compared to mock-inoculated mice (**Fig. 1A and B**). VEEV-TC83 infection (10^5^pfu, i.c.) resulted in a significant increase in CD8+ T-cells as early as day 3 post-infection (**Fig. 1C**). At day 7 post-infection, intranasal (i.n.) inoculation of VEEV-TC83 resulted in disruption of the olfactory bulb tissue architecture in association with a significant increase in CD8+ T-cells (**Fig. 1D and E**). Taken together, these data suggest that the rapid CD8+ T-cell infiltration outpaces the expected kinetics for typical memory T-cell responses and is associated with tissue injury.

**Fig. 1:**
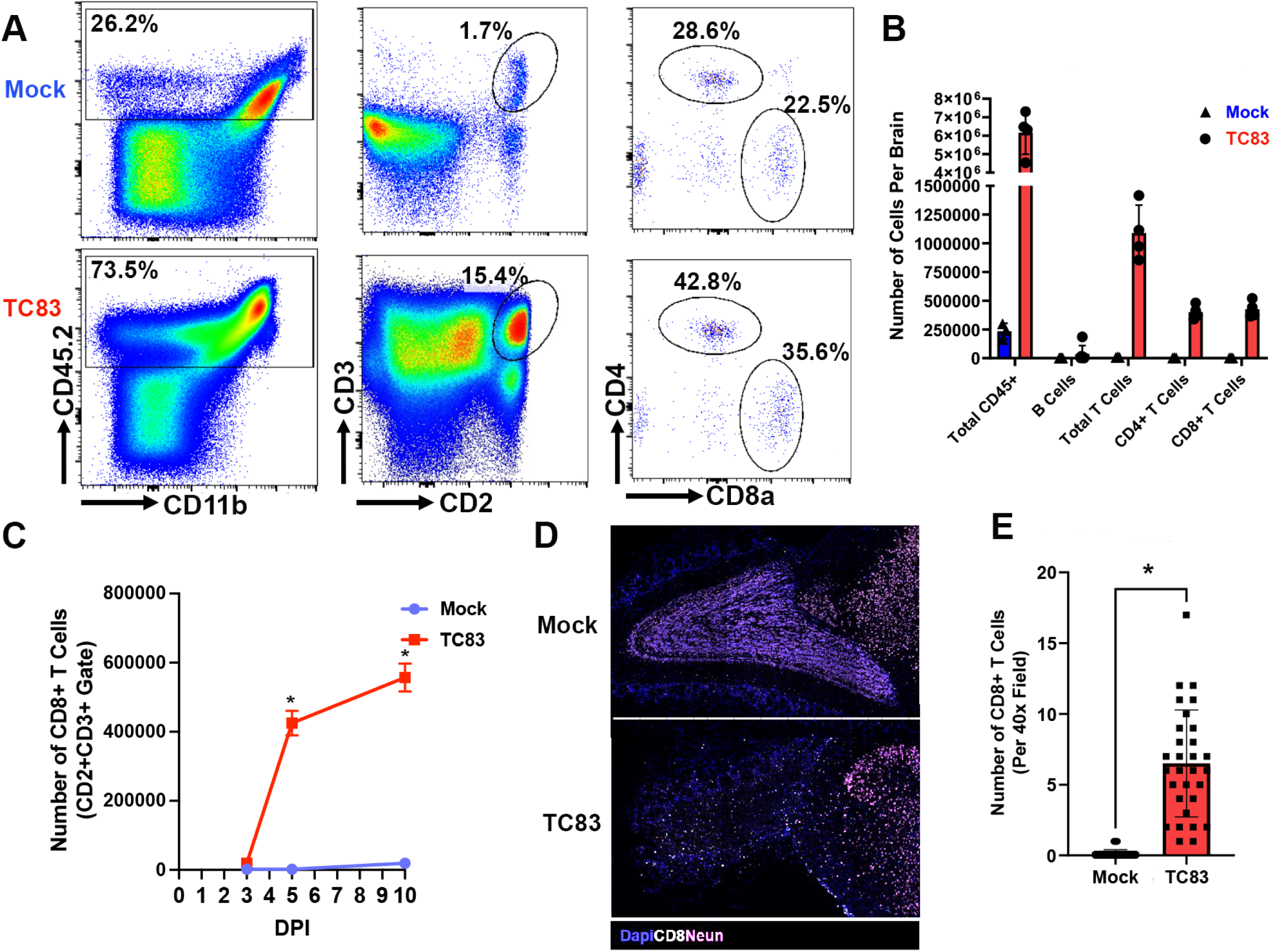
VEEV-TC83 infection induces acute infiltration of T-cells in the olfactory bulb. **A)** Representative flow cytometry plots for identifying CD8+ T cells from the brain (IC, 5dpi, TC83). **B)** Absolute counts of immune cells in brains, (n=4, IC, 5dpi, TC83). **C)** Quantification of flow cytometry analysis identifying total CD8+ T cells in the brains of mice over time, (n=4-6, IC, TC83). **D-E)** IHCp of OBs from mock or infected mice (7dpi, IN, TC83). **D)** Representative images of OBs stained with IHCp (Dapi=blue CD8=white Neun=pink). **E)** Quantification of CD8+ T cells per random 40x high powered field (hpf). (n=30 fields, N=6 OBs, \**p*≤0.05).

### VEEV-TC83 Infection Induces Acute Infiltration of bystander, memory CD8+ T-cells

Next, we determined the phenotype of the CD8+ T-cells that were infiltrating brain tissue at 3 days post-infection. Mice were infected with mock inoculum or TC83 (10^5^pfu, i.c.) as above and brain tissue harvested at 3 days post-infection. We found that TC83 infected mice exhibited a significant increase in CD8+ T-cells expressing markers associated with bystander T cells CD122, CD44, Eomes, CD69, Tbet, CD49d, NKG2D, and CXCR3 (**Fig. 2A and C**). We used the gating of CD44^high^NKG2D+ as a representative strategy for bystander cells, as previous work has demonstrated that these cells are typical bystander CD8+ T-cells.(32) At 10 days post-infection, the population of CD8+ T-cells in VEEV-TC83 inoculated mice exhibited decreased expression of CD122, CD44, Eomes, Tbet, and CD49d suggesting a shift at 10 days from bystander CD8+ T-cells to antigen-specific CD8+ T-cell responses (**Fig. 2B**).

**Fig. 2:**
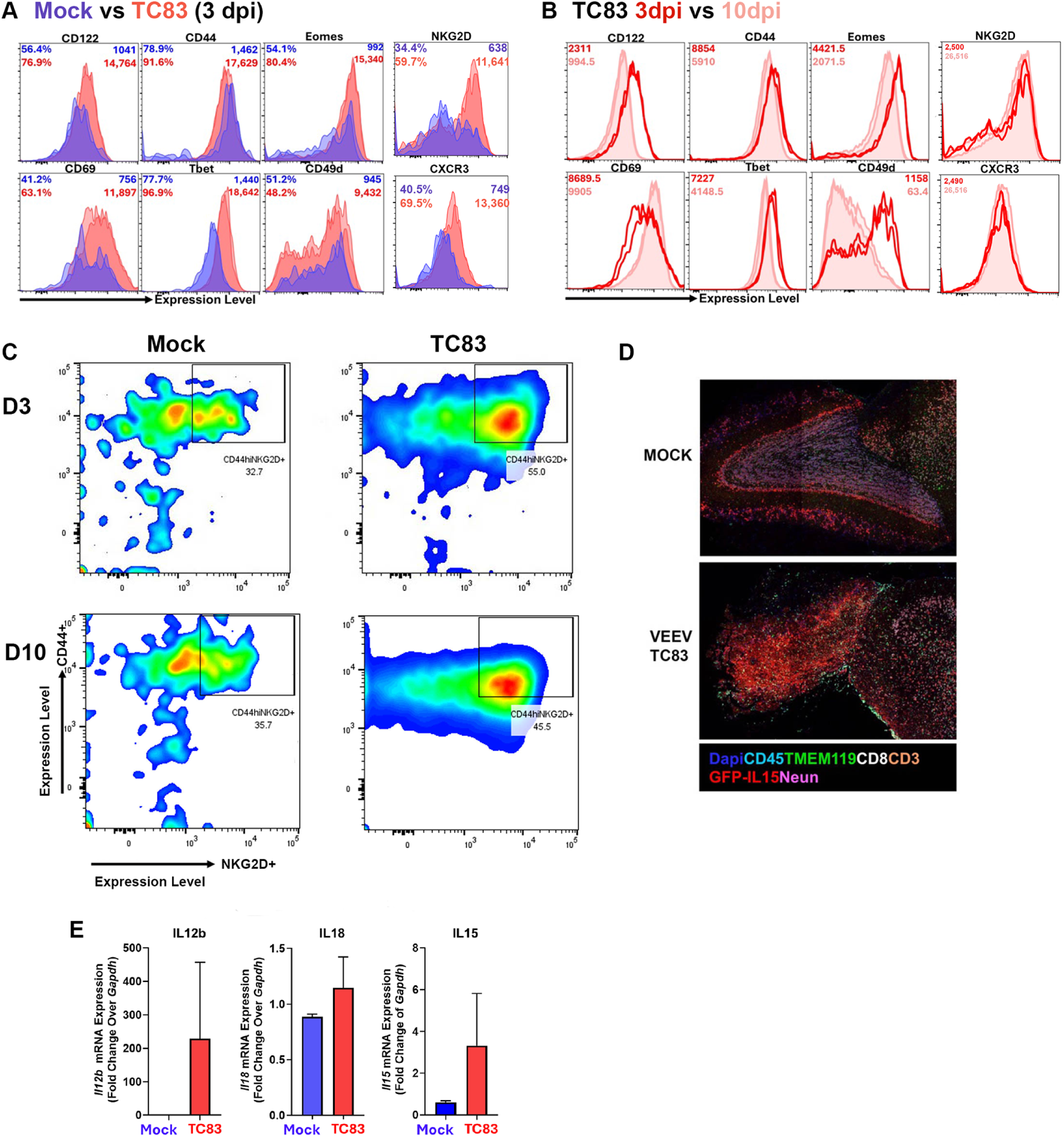
VEEV-TC83 infection induces acute infiltration of bystander, memory CD8+ T-cells. **A)** Representative flow cytometry histograms comparing expression of bystander markers on CD8+ T cells between brains of mock and TC83-infected mice (n=4, 3dpi, IC). **B)** Representative flow cytometry histograms comparing expression of bystander markers on CD8+ T cells between 3dpi and 10dpi (n=4, IC, TC83). **C)** Representative flow contour plots showing CD44 and NKG2D expression on CD8+ T cells at 3dpi (upper panels) and 10dpi (lower panels) (n=4, 3 or 10dpi, IC, TC83) (Gating: Scatter, singlets, live, CD45+, CD2+CD3+, CD8+). **D)** Representative IHCp images from OBs of mock (upper) or infected (lower panel) IL15-reporter mice stained with Dapi=blue, CD45=cyan, TMEM119=green, CD8=white, CD3=orange, GFP-IL15=red, Neun=pink (7dpi, IN, TC83). **e)** qPCR of bystander-associated cytokines (n=4, 5dpi, IC),

Bystander (CD44+CD122+) CD8+ T-cells from infected mice expressed significantly increased IFNγ compared to non-bystander (CD44-CD122-) CD8+ T-cells (**Fig. S2A**).

Additionally, *ex vivo* treatment of immune cells from the brains of TC83-infected mice with IL12 and IL18 resulted in dramatic increases in IFNγ expression (**Fig. S2B**).

We next determined potential cytokine responses found in TC83 infection that are also associated with infiltration of innate, bystander CD8+ T-cells. We searched the NCBI GEO database (GEO Accession GSE91074) and found that IL-15, IL-12b, and IL-18 were associated with TC83 infection in the brain (**Fig. S2C**). Next, we injected GFP-IL15 mice with mock inoculum or VEEV-TC83 (10^5^pfu, i.n.) as above and harvested brain tissue for immunohistochemistry (IHC) analysis at 7 days post-infection. We found that TC83 infection was associated with a significant increase in expression of GFP-IL15 compared to mock inoculated mice (**Fig. 2D**). Consistent with these results, we observed an increase in mRNA expression of IL12b, IL18, and IL15 with TC83 infection at 5dpi in WT mice (**Fig. 2E**). These data show that infiltration of innate bystander CD8+ T-cells occurs at early time points and is associated with IL-15 expression in the brain.

### Microglia and Macrophage-dependent IL15 production supports bystander CD8+ T-cell recruitment and cytotoxicity

Next, we determined the source of IL-15 production and the role of IL-15 in activation of infiltrating CD8+ T-cells in the brain. Wild-type mice were inoculated with mock inoculum or TC83 (10^5^pfu, i.n.) and brain tissue was harvested at day 5 post-infection. We found that TC83-inoculated mice exhibited increased numbers of microglia (CD45^+^, CD11b^mid^) and increased numbers of infiltrating macrophages (CD45^high^, CD11b^high^) compared to mock-inoculated mice (**Fig. 3A**). Gating on these populations, we found that both resident microglia and infiltrating macrophages express high levels of IL-15 with macrophages expressing the highest levels of IL-15 based on the IL-15-GFP reporter expression (**Fig. 3A**).

**Fig. 3:**
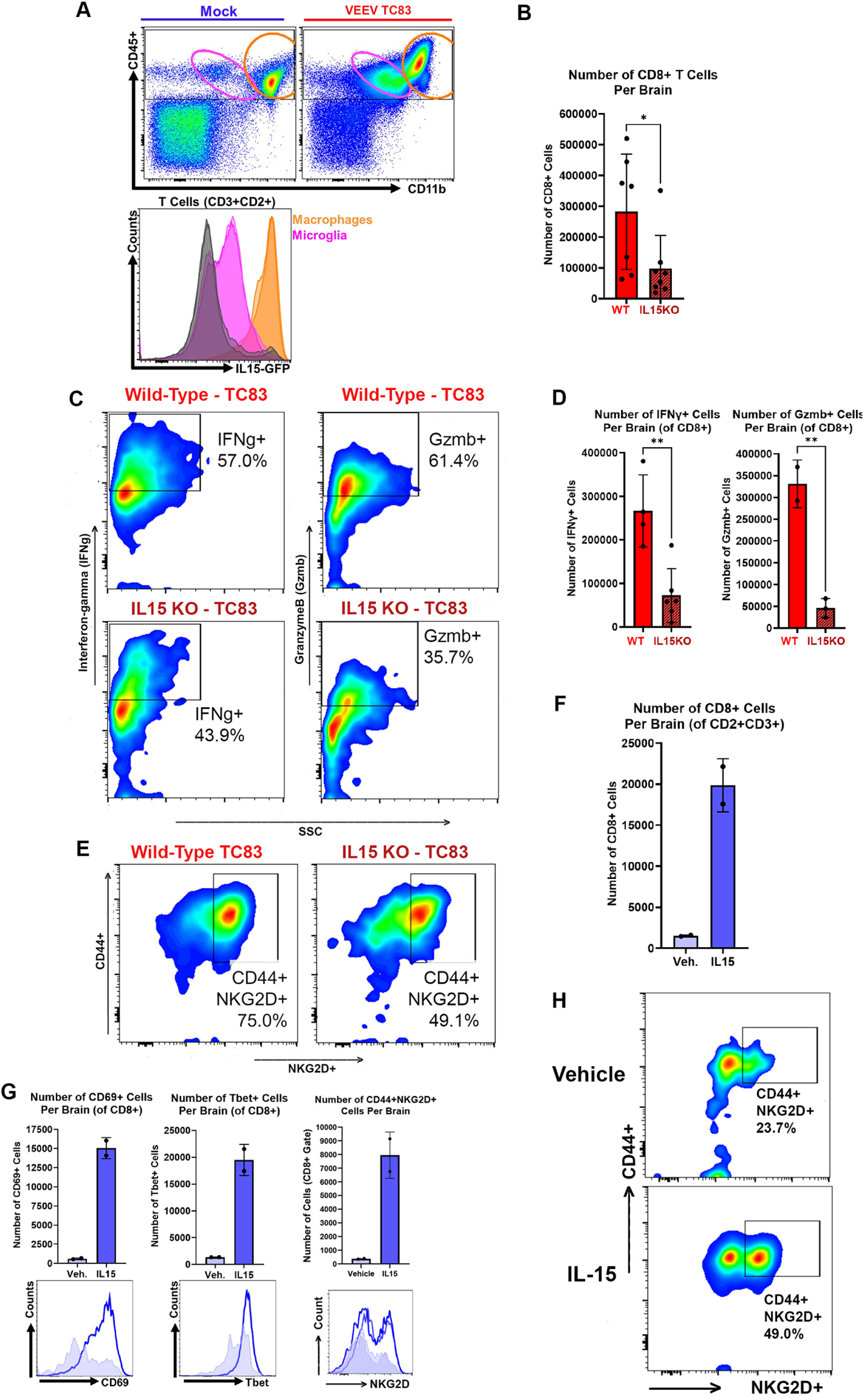
Microglia and macrophage-dependent IL15 production supports CD8+ T-cell recruitment and cytotoxicity. **A)** Upper panel: Representative flow plots of macrophage (CD11bhiCD45+, orange) and microglial (CD11b^mid^CD45+, pink) populations. Lower panel: histogram plots of IL15-GFP expression in macrophages (orange), microglia (pink) and T cells (black). (Gating: scatter, singlets, live; n=4, IC). **B)** Quantification of CD8+ T cells in the brains of infected WT or IL15ko mice (n=7, IC). **C)** Representative flow plots of cytokine and surface marker expression from CD8+ T cells from WT (upper panels) or IL15ko (lower panels) mice, all infected with TC83. (n=6, IC). **D)** Quantification of cytokine expression in CD8+ T cells of WT or IL15ko mice (IFNγ: n=4; Gzmb: n=3, all IC). **E)** Flow cytometric analysis of CD8+ T-cells from brains of WT and IL15KO mice indicating CD44^high^NKG2D+ cells (n=6, IC). **F)** Mice were treated with vehicle or rIL15 (5ug) at day 0 and again 48hrs later. Mice were harvested 48hrs post first dose (n=4, IC). **G)** Quantification of protein expression from flow cytometric analysis (top panels) of CD8+ T cells with representative flow histograms (lower panels) from vehicle or rIL15 (5ug) treated mice (vehicle=light blue, rIL15=dark blue; IC, n=4). **H)** Representative flow plots of CD44^high^NKG2D+ CD8+ T-cells in vehicle or rIL15 treated mice (n=4, IC).

To define the role of IL-15 in supporting both CNS infiltration and activation of bystander CD8+ T-cells, we inoculated IL-15 knockout (KO) mice with mock inoculum or VEEV-TC83 (10^5^pfu, i.c.) and harvested brain tissue for spectral flow cytometry at day 5 post-infection. We found that IL-15 KO mice had a significant reduction (66%) of CD8+ bystander T-cells (**Fig. 3B**). Moreover, TC83 inoculated IL-15 KO mice also exhibited a significant reduction in activation of CD8+ T-cells based on interferon-gamma and granzyme B production (**Fig. 3C and D**). In addition to decreased activation markers, the CD8+ T-cells in the brains of infected IL15KO mice exhibited decreased surface marker expression associated with the bystander phenotype (**Fig. 3E**).

Next, we determined if IL-15 alone was able to recruit CD8+ bystander T-cells. Wild-type mice were inoculated with IL-15 (5ug, i.c. inoculation) at day 0 and day 2, and brain tissue harvested at day 4 post-first-injection for spectral flow cytometry analysis. We found that, compared to vehicle control, IL-15 inoculation induced a significant increase in numbers of infiltrating CD8+ T-cells that were CD69+ and Tbet+ (**Fig. 3F and G**). Additionally, IL15 injection was sufficient to increase the expression of bystander-associated surface markers CD44 and NKG2D (**Fig. 3G and H; Fig. S3**). We also observed increased expression of other bystander-associated markers with Poly I:C treatment and IL15 treatment including CD44+, CD122+, NKG2D+, Tbet+, CD49d+, Eomes+, and CD69+ (**Fig. S3**). These results are consistent with activated bystander T-cell populations following activation by RNA detection and IL-15 activation, respectively.

### Virus-induced cytotoxic, bystander CD8+ T-Cells in the Brain are activated independent of the T-cell Receptor

Next, we determined if bystander CD8+ T-cell populations in the brain were activated independent of the T-cell receptor (TCR). For these experiments, we utilized OT-I mice that carry a transgene containing the rearranged mouse Tcra-V2 and Tcrb-V5 genes encoding a TCR that recognizes a chicken ovalbumin peptide (257–264) bound to MHC class I. In these mice, nearly all the T-cells are ovalbumin-specific CD8+ T-cells and, thus, should not have an antigen-specific response to any other antigen. Wild type and OT-I mice were injected with mock or TC83 inoculum (10^5^pfu, i.n.) and brain tissue was harvested at day 5 post-infection for spectral flow cytometry analysis. We found that OT-I mice exhibited significantly increased infiltration of CD8+, Tbet+ T-cells following VEEV-TC83 infection compared to mock-inoculated mice and CD8+ T-cell infiltration was similar in OT-I mice compared to WT mice (**Fig. 4A**).

**Fig. 4:**
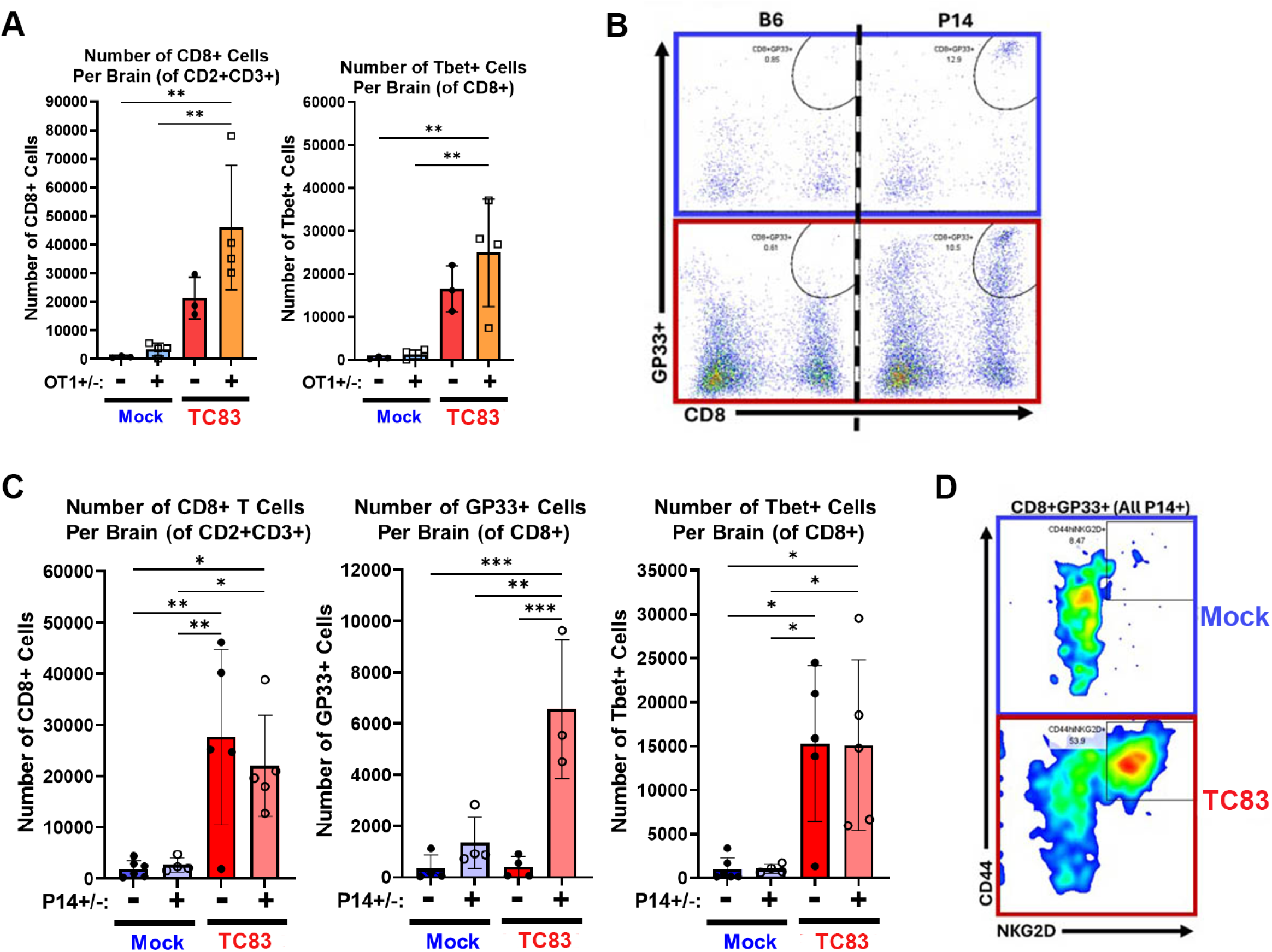
Virus-induced cytotoxic bystander CD8+ T-cells in the brain are activated independent of the T-cell receptor. **A)** Quantification of flow cytometric analysis of CD8+ T cell numbers from mock or infected WT or OT1 TCR transgenic mice (n=3-4, IC, TC83). **B)** Representative flow plots of GP33 tetramer+CD8+ T cells from mock (upper panels) or TC83 infected (lower panels) WT or P14 TCR transgenic mice (n=5, IC). **C)** Left panel: Quantification of total CD8+ T cell numbers per brain. Middle panel: Quantification of GP33-specific CD8+ T-cells from B. Right panel: Quantification of Tbet+ CD8+ T-cells per brain. (IC, TC83, n=4-5, *p*≤0.5, one-way ANOVA with Multiple Comparisons post analysis). **D)** Representative flow plots of CD44^high^NKG2D+ CD8+ T cells from P14 TCR transgenic mice mock (upper panel) or TC83 infected (lower panel) (n=4-5, i.c., TC83).

To further evaluate antigen-specific CD8+ T-cell responses, we used P14 TCR transgenic mice that express a TCR specific for lymphocytic choriomeningitis virus (LCMV) due to the availability of tetramers and other assays to define TCR-specific responses. The P14 TCR recognizes the LCMV gp33 peptide (33–41); furthermore, this line is crossed to a TCRα KO background to prevent development of endogenous TCR αβ T cells expressing any other TCR and should therefore not respond to any other antigen. We infected WT (P14-) or P14+ mice i.n. with TC83 and then analyzed the brain for tetramer+ CD8+ T-cells after 5 days following TC83 infection. Despite TC83 lacking LCMV antigens, there was a significant increase in total CD8+ T-cell and GP33-specific (the dominant LCMV epitope for MHCI) CD8+ T-cell infiltration into the brain at 5dpi (**Fig. 4B and C**). Moreover, these GP33-specific CD8+ T-cells expressed increased levels of Tbet were largely CD44^high^NKG2D^+^, indicating activation and bystander phenotype (**Fig. 4D**). Consistent with these results, TC83-infected P14 mice had similar numbers of CNS-infiltrating bystander CD8+ T-cells to TC83-infected WT or P14-mice as defined by Tbet+, CD122+, CD44+NKG2D+, CD69+, Eomes+, NKG2D+ (**Fig. S4**). CD8+ T-cells from P14 mice are not capable of antigen-driven activation by any antigen other than gp33, yet CD8+ T-cells in P14+ mice following TC83 infection are activated and capable of producing IFNγ indicating mechanisms of activation are independent of the TCR.

Next, we determined if infiltrating bystander CD8+ T-cells in the brain exhibit evidence of cytotoxicity. Wild type mice were inoculated with mock or VEEV TC83 inoculum (10^5^pfu, i.n.) and brain tissue harvested at day 5 post-infection for analysis as above. We found that CD8+ T-cells exhibit a significant increase in markers of cytotoxicity including CD107a, interferon-gamma, and granzyme B (*p*≤0.05, student’s t-test, **Fig. 5**, **Fig. S2A**). Moreover, CD8+ T-cells from TC83-infected mice which express bystander markers (CD44+CD122+) produce significantly more IFNγ than CD8+ T-cells from the same mice not expressing bystander markers (CD44-CD122-; **Fig. S2A**).

**Fig. 5:**
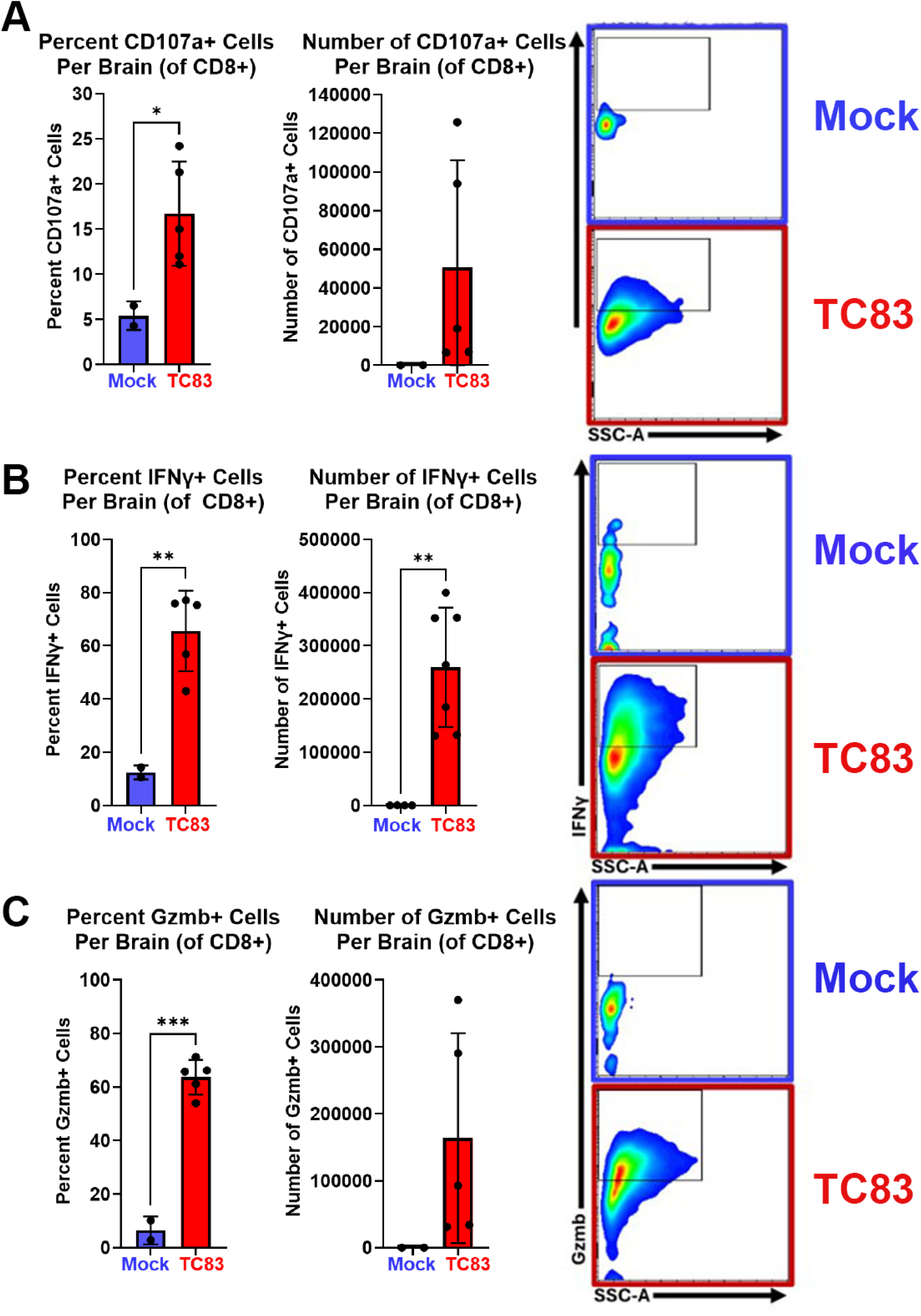
Bystander CD8+ T cells from the brains of TC83-infected mice exhibit cytotoxic characteristics. Left panels are quantification, right panels are representative flow plots. **A)** Flow cytometric analysis of CD107a expression in CD8+ T cells (n=5, IC, TC83). **B)** Flow cytometric analysis of IFNγ production in CD8+ T cells (n=6, IC, TC83). **C)** Flow cytometric analysis of Gzmb expression in CD8+ T cells (n=5, IC, TC83). \**p*≤0.05, Student’s t-test.

These data show that infiltrating bystander CD8+ T-cells exhibit cytotoxic phenotypes that may contribute to tissue injury.

### Bystander CD8+ T-cell infiltration and activation is dependent on type 1 interferon

We next determined if acute infiltration of bystander CD8+ T-cells in the brain was stimulated by antigen detection and interferon production. Wild-type mice (6-10 weeks of age) were inoculated by intracranial injection with phosphate buffered saline (PBS, 10ul), poly I:C (10ug/10ul), interferon-alpha2 (2x10^4^ Units/10ul), or interferon-beta (1x10^4^/10ul) at day 0 and again day 2 post-treatment. At day 4 following poly I:C treatment, we found that gene expression in brain tissue of IL-12b and IL-15 were significantly increased compared to control treated mice in association with increased expression of CD8+, CD44+, and NKG2D+ T-cells in the brain (**Fig. 6A and B**). Next, we determined the role of type 1 interferon in recruitment of bystander CD8+ T-cells into brain tissue in the same treatment groups. At day 4 post-treatment with interferon-alpha, we found increased gene expression of IL12b and IL-15 compared to brain tissue treated with control (**Fig. 6C**). Following treatment with interferon-alpha, we also found significantly increased infiltration of CD8+, CD44+, and NKG2D+ T-cells in brain tissue (**Fig. 6D**). We also found that mice treated with type 1 interferon exhibit increased numbers of CD8+, Tbet+, CD69+ T-cells compared to PBS treated brain tissue (**Fig. 6E and F**). These data show that interferon contributes to expression in the brain of IL-15, IL-12, and infiltration of bystander CD8+ T-cells.

**Fig. 6:**
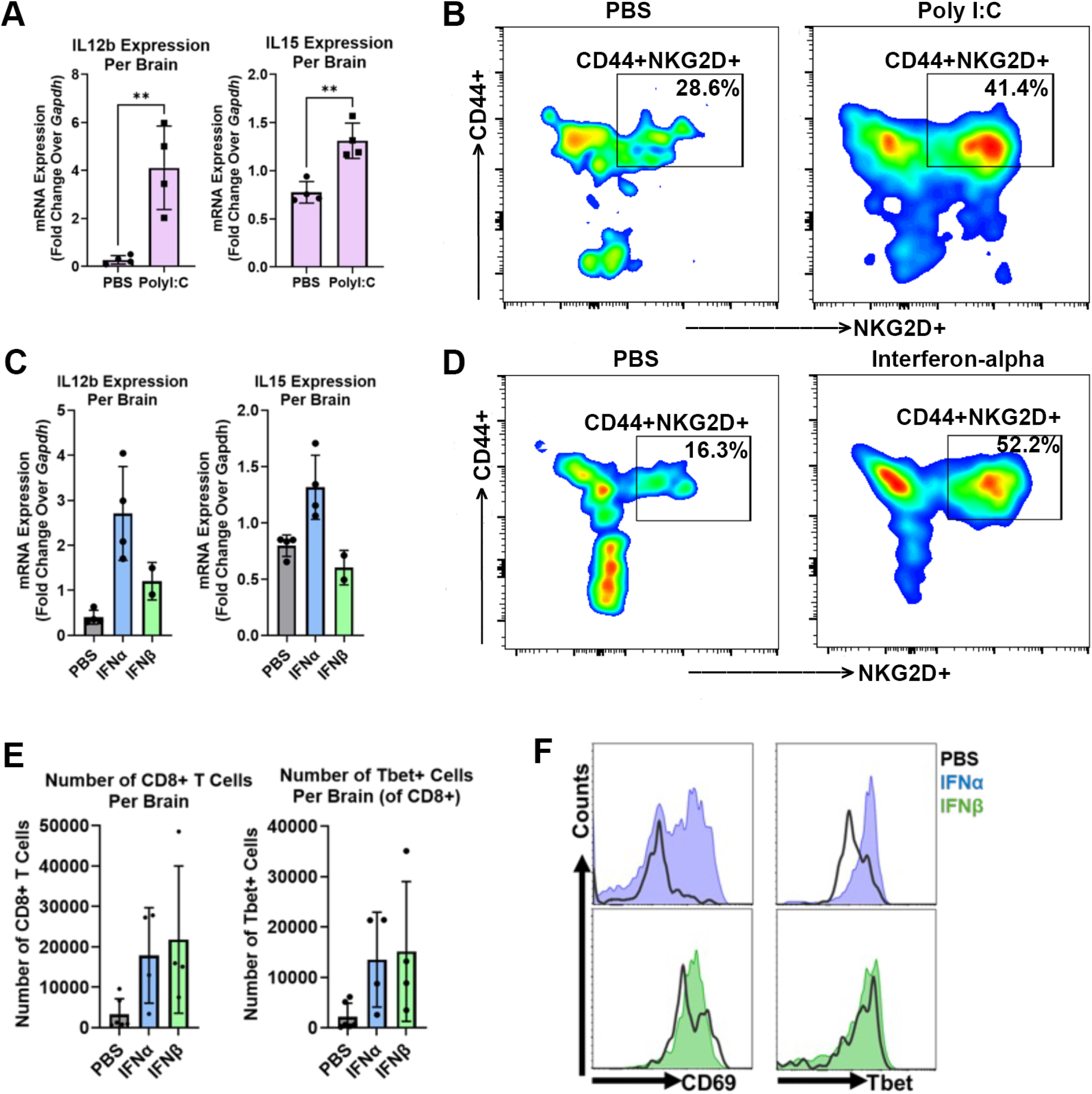
Bystander CD8+ T cell infiltration and activation is dependent on type 1 interferon. Mice were treated IC with vehicle (PBS) or PolyI:C, IFNa, or IFNb at 0hrs and again at 48hrs. Mice were harvested 4 days after first injection. **A)** Gene expression from mice treated with vehicle or PolyI:C (n=4). **B)** Representative flow plots of CD44hiNKG2D+ CD8+ T cells from PBS or PolyI:C treated mice. **C)** Gene expression from IFNα or IFNβ treated mice (n=4). **D)** Representative flow plots of CD44hiNKG2D+ CD8+ T cells from PBS, IFNα, or IFNβ treated mice. **E)** Quantification of CD8+ T-cells (left panel) and Tbet+CD8+ T cells (right panel) per brain from PBS, IFNα, or IFNβ treated mice. **F)** Representative flow cytometry histograms of activation markers on CD8+ T-cells from PBS (black outline), IFNα (blue), or IFNβ (green) treated mice (n=4). \*\**p≤*0.05, student’s t-test or one-way ANOVA with multiple comparisons post analysis).

To demonstrate the importance of type 1 interferon for bystander CD8+ T-cell recruitment and activation, we infected type 1 interferon receptor knockout (IFNARKO) mice i.c. with 10^5^PFU TC83. Mice were sacrificed at 2 days post infection because IFNARKO had become moribund. Despite the early timepoint of harvest due to moribund mice, we observed bystander phenotypes of CD8+ T-cells infiltrating the brains of WT mice (**Fig. S5**). CD8+ T-cells infiltrating the brains of TC83-infected IFNARKO mice expressed lower levels of bystander markers (CD44^high^NKG2D+, CD69+) suggesting that type 1 IFN is important for bystander CD8+ T-cell recruitment and activation (**Fig. S5**).

To investigate whether bystander CD8+ T-cell recruitment and activation is a general response to viral infection, we infected mice with LCMV (200pfu, or 500pfu, i.c.) and 5 days post infection evaluated their CNS-infiltrating CD8+ T-cells. Similar to TC83 infection, infiltration of CD8+ T-cells from the brains of LCMV-infected mice also occurred early (5dpi) (**Fig. S6**). Additionally, the CNS-infiltrating CD8+ T-cells exhibit bystander marker expression (CD44^high^NKG2D+, NKG2D, CD69, Tbet, CD49d, Eomes; **Fig. S6**). In addition to the type 1 interferon data, these data strongly suggest that bystander CD8+ T-cell infiltration into the brain is dependent on type 1 interferon production and nonspecific pathogen response to viral infection of the brain.

### Bystander CD8+ T-cells contribute to CNS injury following virus infection

We next determined the role of bystander CD8+ T-cell infiltration in virus control and tissue injury in the CNS. Wild-type and RAG1 knockout (KO) mice were inoculated with VEEV-TC83 (10^5^pfu, i.n) and brain tissue analyzed at 5 days post-infection for virus replication. We found that there was no significant difference in TC83 genome copies in RAG1 KO brains compared to WT mice at this early time point (**Fig. 7A**). Next, we treated WT mice with an antibody specific for CD8 (250ug i.p.) or isotype control to deplete CD8+ T-cells. At 7 days post-treatment, mice were inoculated TC83 (10^5^pfu, i.n) and brain tissue analyzed at 5 days post-infection and CD8+ T-cell depletion verified in the CNS via flow cytometry (**Fig. S7**). We found that CD8+ T-cell depletion did not significantly alter TC83 genome copies in the brain tissue at 5 days post-infection compared to isotype control treated mice (**Fig. 7B**). Upon analysis of immunofluorescent histology of brain sections from treated mice, we found that mice treated with antibody depletion of CD8+ T-cells exhibited significantly decreased expression of cleaved-caspase 3 in NeuN+ cells and GFAP+ cells (**Fig. 7C and D**). We also found decreased expression of phosphorylated stat1 in GFAP+ cells in VEEV-TC83 inoculated brain tissue from mice treated with CD8 depletion antibody (**Fig. 7C and E**). Together, these data show that acute infiltration of bystander CD8+ T-cells does not alter early virus replication while contributing to increased caspase-3 activity, especially in neurons and astrocytes.

**Fig. 7:**
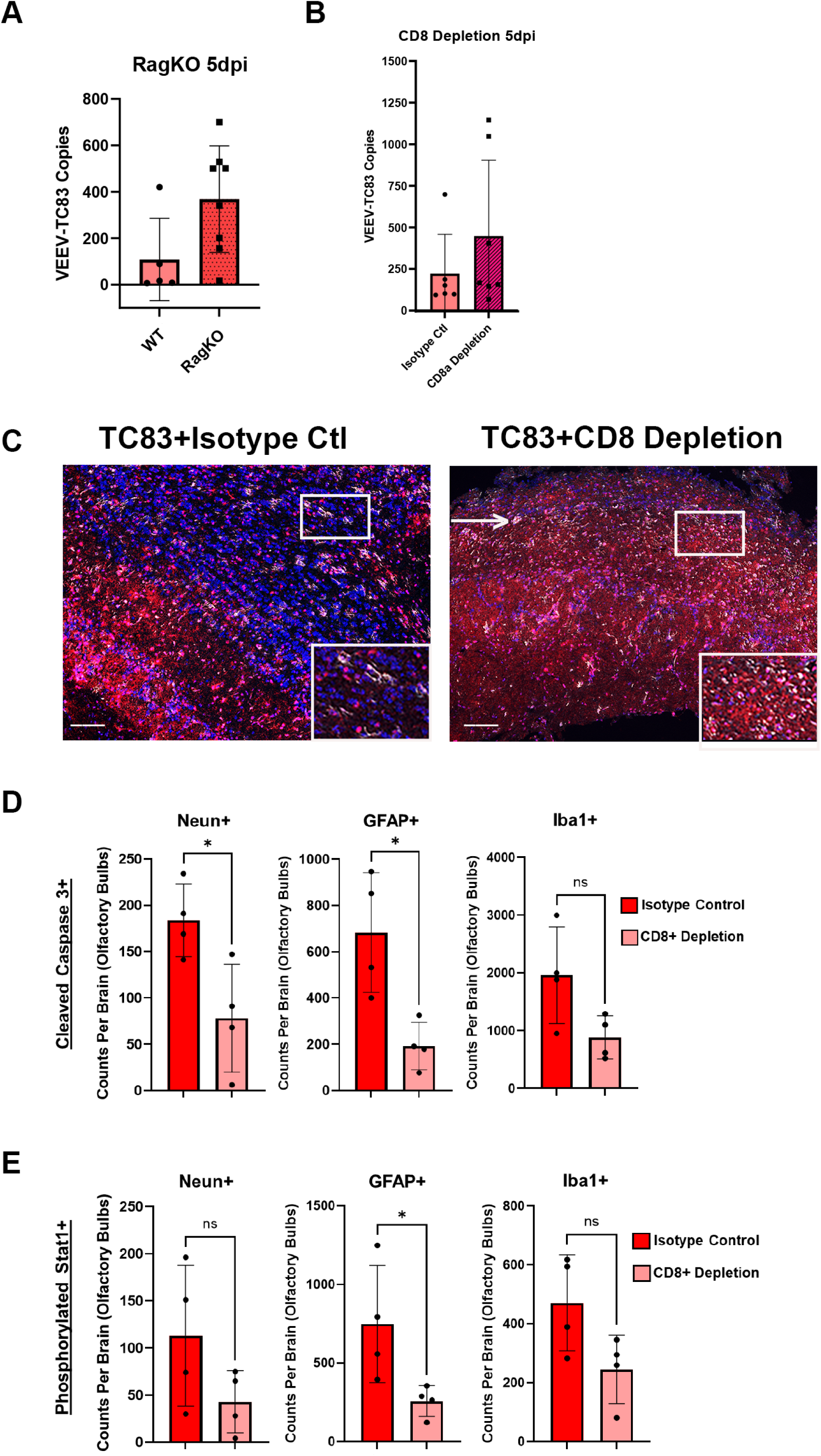
Bystander CD8+ T cells contribute to CNS injury following virus infection. **A)** Rag-/-mice and WT mice were i.c. infected with TC83 and viral genome copies in the brain were measured with qPCR (5dpi, n=8). **B)** At the time of ic infection, WT mice were injected ip with either isotype control antibody or CD8-depletion antibody. Viral genome copies were measured with qPCR (5dpi, TC83, n=8). **C)** Mice were infected IN with TC83 and either injected IP with isotype control antibody or CD8-depletion antibody. At 5dpi, brains were harvested and processed for IHCp. Representative images of the OBs of infected mice (Dapi=blue, Neu=pink, cl-caspase3=white). **D)** Cleaved caspase 3 was quantified with IHCp of individual OBs in neurons (Neun+), astroglia (GFAP+), and microglia (Iba1+) (n=4, *p*≤0.05 Student’s t test). **E)** Phosphorylated Stat1 was quantified with IHCp of individual OBs in neurons (Neun+), astrocytes (GFAP+), and microglia (Iba1+) (n=4, \**p*≤0.05 Student’s t test).

## Discussion

Our results provide an important mechanism by which innate, bystander cytotoxic CD8+ T-cell populations in the brain are recruited and activated independent of antigen-specific T-cell receptor (TCR) stimulation. We found that both type I interferon and IL-15 responses by myeloid cell populations of resident microglia and infiltrating macrophages support recruitment and activation of bystander CD8+ T-cells in the brain. Previous work with Zika virus infection in a type I interferon knockout mouse model has shown that bystander CD8+ T-cells infiltrate the brain and contribute to neuropathology.(33) Since these studies were completed in type 1 interferon receptor knockout mice, the role of interferon-dependent responses, relevant cytokine responses, and T-cell antigen-specific responses were not completely evaluated. Prior work with a reovirus model of infection has shown that activation of bystander CD8+ T-cells in the spleen occurs as early as 24 hours post-infection and is dependent on type 1 interferon.(34) However, the mechanisms that induces infiltration and activation of bystander CD8+ T-cells in the brain were not previously understood.

We found that bystander CD8+ T-cells infiltrate the CNS in part due to type 1 interferon responses in the brain and IL-15 production from microglia and infiltrating macrophages following virus infection. Injection of type 1 interferon or poly I:C was sufficient to induce acute infiltration of bystander CD8+ T-cells into the brain. Type 1 interferon is known to induce IL-15 and related NK-cell and T-cell activating cytokines in antigen presenting cells such as dendritic cells.(35, 36) Additionally, IL-15 is activated in astrocytes and microglia during acute injury in the CNS from intracerebral hemorrhage or acute neuroinflammation using lipopolysaccharide in mice.(37, 38) In IL-15 knockout mice, we found significant reductions in the infiltration and activation of bystander CD8+ T-cells in the brain following virus challenge. Our studies provide an important new mechanism by which bystander CD8+ T-cells are recruited to the CNS.

Similar to prior studies,(33) our data also show that bystander CD8+ T-cells do not contribute to significant control of virus replication at early time points; however, depletion of CD8+ T-cells at early time points results in decreased injury in the CNS with a decrease in cleaved-caspase 3 in neurons and astrocytes in brain tissue. These data provide a model by which stimulation of acute type I interferon dependent expression of IL-15 results in early infiltration of bystander CD8+ T-cells that contribute to CNS injury and neuropathology at early time points.

Importantly, our work also evaluates the role of virus infection in olfactory pathways in the recruitment and neuropathology associated with bystander CD8+ T-cells. A recent study of cognitive deficits in patients with COVID-19 found objectively measurable cognitive deficits that may persist for a year or more after infection.(39) Additionally, respiratory viruses such as SARS-CoV2 are known to cause inflammation in olfactory pathways resulting in the potential for neuroinflammation and long-term cognitive outcomes.(40) However, the mechanisms linking viral exposure in olfactory pathways with neuropathologic responses are unclear. Our data suggest that olfactory virus infection may be sufficient to trigger type 1 interferon dependent responses in the olfactory bulb and brain tissue resulting in activation of IL-15 dependent activation of neuropathologic bystander CD8+ T-cells. Future studies evaluating the role of respiratory viruses and olfactory pathway stimulation of bystander T-cell responses in the brain will be important to understand the mechanisms linking acute respiratory virus infection and cognitive outcomes in infections like COVID-19.

Recent studies have shown that individuals infected with Epstein-Barr virus (EBV) exhibit an increased risk of developing multiple sclerosis.(41) These patients exhibit increased numbers of T-cells that exhibit antigen specificity for EBV peptides that overlap with self-antigens in the CNS. However, the mechanisms by which memory CD8+ T-cells are stimulated to enter the CNS are not clear. EBV-specific CD8+ T cells selectively infiltrate the brain in multiple sclerosis and interact with virus-infected cells, and peripheral HIV-p18 specific, bystander CD8+ T-cells initially infiltrate brain tissue following challenge with neuroinvasive mouse coronavirus infection.(42, 43) Our data provide a potential mechanism by which pathological memory T-cells may enter the CNS and engage potential autoreactive self-antigens in the CNS resulting in initiation of autoimmune inflammatory responses. Further studies are needed to evaluate the role of olfactory nerve inflammatory responses and CNS myeloid interferon and IL-15 dependent responses in the recruitment of peripheral, autoreactive memory T-cells.

In summary, we propose a novel model by which olfactory virus exposure induces interferon-dependent, IL-15 expression in microglia and infiltrating macrophages in the CNS resulting in infiltration and activation of neuropathologic bystander CD8+ T-cells. These results provide a mechanism by which T-cells may gain access to the CNS during acute inflammatory signals in the olfactory track and cause CNS injury. Further studies evaluating these mechanisms may provide novel therapeutic targets for immune modulation to prevent and treat T-cell dependent neuropathology in the brain.

## Acknowledgements

JDB and LJB are supported in part by NIH/NINDS R01 NS123431. Additionally, JDB is supported by NIH/NIAID R01 AI153724. We thank Ross Kedl for generously sharing his IL15-GFP reporter mice. We also thank Mark D’Antonio from Raul Torres’ lab for generously sharing LCMV reagents and knowledge. We want to thank Camille Merrick and Maddeline Slater for providing technical expertise in support of the in vivo experiments.

## Author Contributions

JB led the experimental studies, performed the experiments, analyzed data, and wrote the manuscript. AR, DH, and BM assisted with experiments. JDB helped to conceive of the project, obtained funding, oversaw the experimental work, analyzed data, synthesized data and figures, and participated in writing the manuscript. LJB helped to conceive of the project, obtained funding, oversaw the experimental work, and edited the manuscript.

## Methods

### Mouse Experimental Procedures

All mouse experimental protocols and procedures were approved by the local IACUC. Mice were age matched, and sex matched between the ages of 6-10 weeks old. Mice were anesthetized with isoflurane and depth of sedation tested with response to foot pinch. Once anesthesia was induced, mice were inoculated per protocol by intracranial (i.c.), intranasal (i.n.), or intraperitoneal (i.p.) injection. P14 TCR transgenic mice used for in vivo studies were a gift from Susan Kaech (Yale). IL-15 knockout and IL-15 reporter mice were a gift from Ross Kedl (University of Colorado Anschutz Medical Campus). Wild-type C57/B6 mice were purchased from Jackson labs (Maine). See table 1 for list of mouse strains.

**Table 1.**
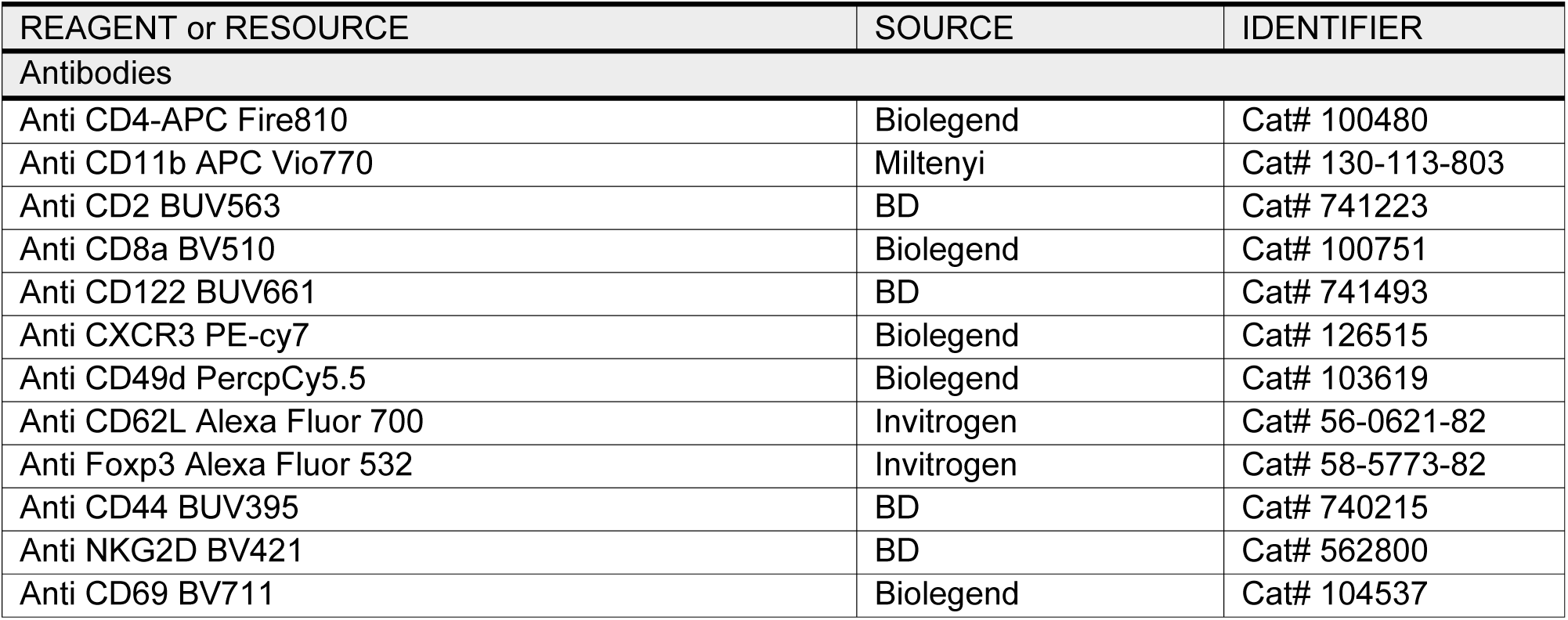
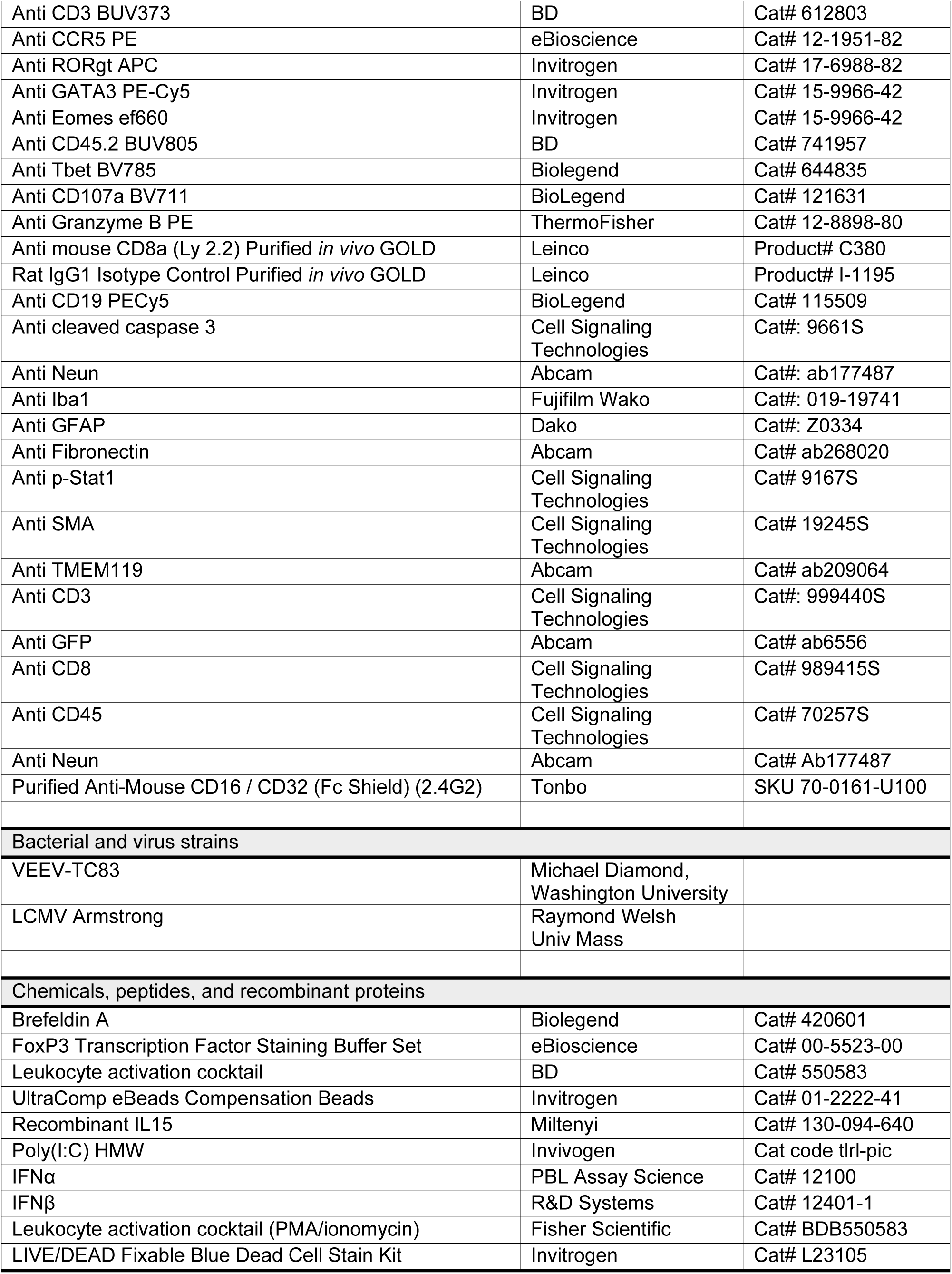
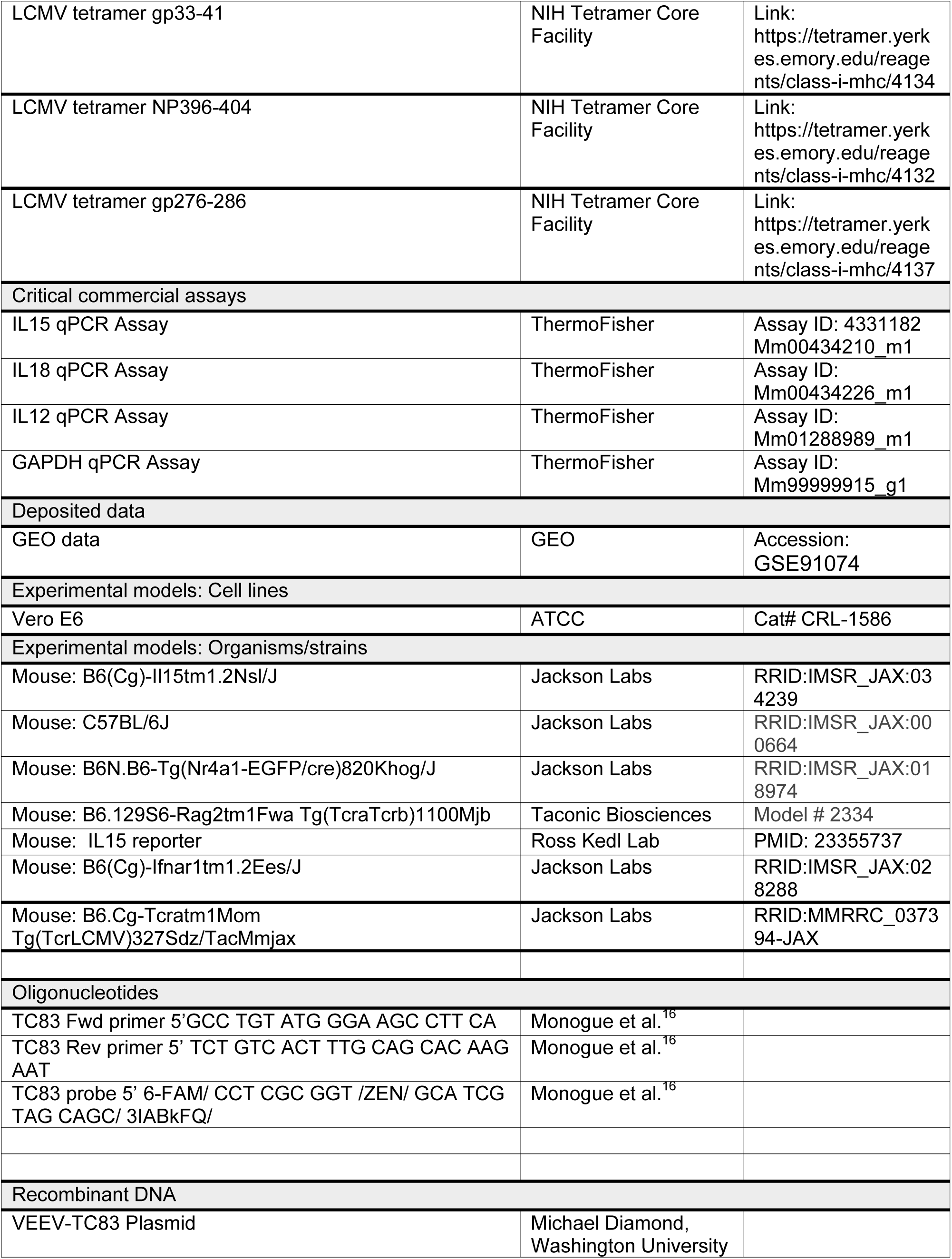
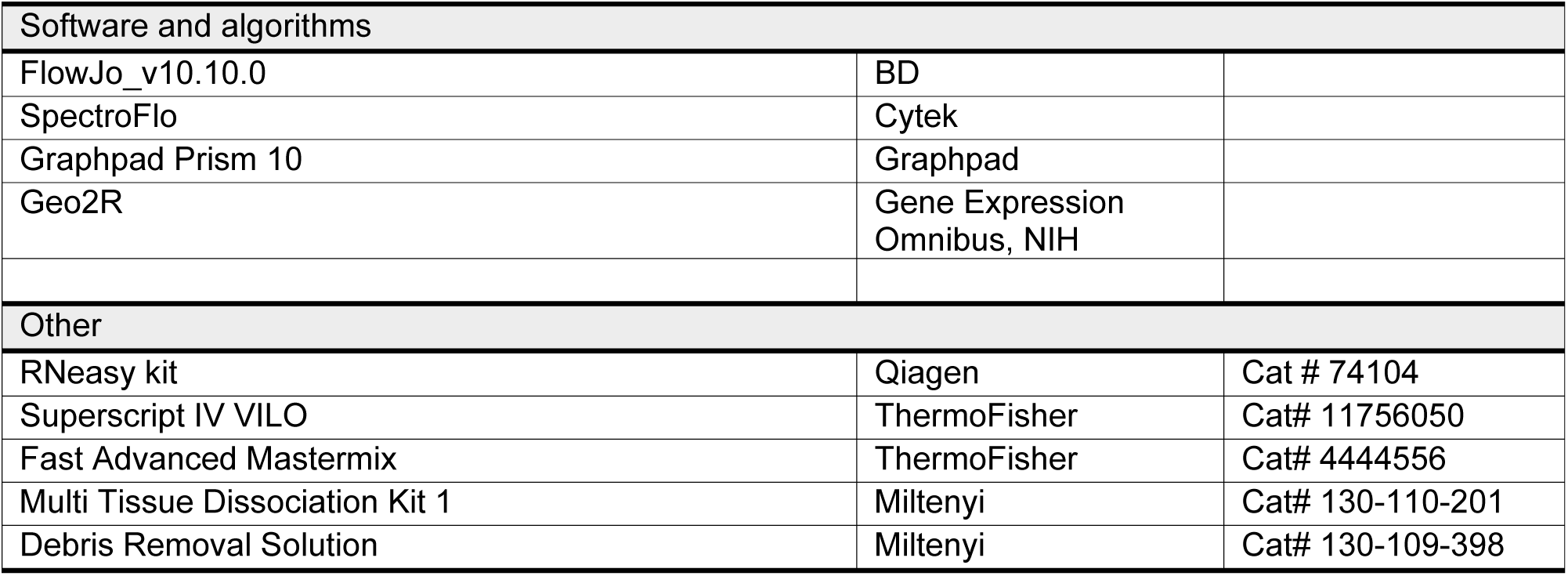

### Flow cytometry

Brain tissue was processed using the Multi Tissue Dissociation Kit 1 following the manufacturer’s protocol (Miltenyi). Cells were counted using a hemacytometer and plated for flow cytometry staining. Spleen tissue was smashed through a 40um filter (Greiner-bio) and red blood cells were lysed with ACK lysis buffer (Gibco) per manufacturer’s protocol. Cells were stained with LIVE/DEAD blue fixable stain kit (Invitrogen) and Fc block (Tonbo) for ten minutes followed by cell surface staining in flow buffer [1xPBS (Fisherscientific) with 1% FBS and 1mM EDTA (Millipore Sigma)] for 20 minutes at room temperature in the dark. Intracellular staining was completed using the FoxP3 Transcription Factor Staining Set (eBiosciences) for 30 minutes at 4°C. Staining for CD107a was completed overnight *ex vivo* prior to any other staining. LCMV-specific tetramers for the three dominant MHC1 epitopes were acquired from the NIH Tetramer

Core Facility at Emory University. For cytokine staining, cells were isolated as above and cultured *ex vivo* for 4hrs with BFA (BioLegend) alone or leukocyte activation cocktail containing BFA (PMA/Ionomycin Fisher Scientific).

All antibodies were titrated to appropriate dilutions for staining brain tissue (Table 1). Flow cytometry analysis was conducted using the Cytek Aurora instrument with the SpectroFlo software (Cytek). Unmixing was completed using single stained UltraComp eBeads (Invitrogen). Data analysis of FCS files was performed with FlowJo_v10.10.0.

### Viruses

VEEV-TC83 (gift from Michael Diamond, Washington University) was grown in Vero E6 cells (ATCC). Virus titer was measured using plaque assays on Vero E6 cells. LCMV Armstrong stocks were propagated in baby hamster kidney 21 cells and generously provided by Dr. Raymond M Welsh and amplified in the Berg laboratory.

### Antibody CD8+ T-Cell depletion

To deplete CD8+ T-cells, mice were injected i.p. with 250ug of isotype control (Leinco, clone GL113) or CD8a depletion antibody (Leinco, clone 2.43) at the time of infection with TC83. Successful CD8+ T-cell depletion was verified after harvest using flow cytometry.

### RT-qPCR

Minced brain tissue was placed in 1 mL of RLT buffer from the RNeasy RNA isolation kit (Qiagen) and stored at -20°C until ready for processing. Tissue in RLT buffer was heated to 65°C for 10min to ensure virus inactivation. Once thawed, tissue was homogenized using the Bead Bug (Benchmark) for 60 seconds at 400RPM. 350ul of homogenized tissue in RLT was carried forward for RNA isolation with the RNeasy kit (Qiagen). RNA quality and quantity was measured with the NanoDrop spectrophotometer (Invitrogen). Relative gene expression to GAPDH and fold change over GAPDH were calculated as previously described.(44) CDNA was synthesized from RNA using the SuperScript IV VILO mastermix (Thermo Fisher Scientific).

Gene expression was quantified using 20x TaqMan gene expression assays and the Fast Advanced Mastermix (Thermo Fisher Scientific). qPCR was performed with the Quant Studio 3 thermocycler (Applied Biosystems). VEEV-TC83 genome copies were quantified using primers and probes to nsp1 with the Fast Advanced Mastermix (Thermo Fisher Scientific)^16^. See table 1 for list of primers and probes. To calculate genome copies, Ct values were compared to a standard curve made from diluted plasmid containing the TC83 genome.

### Imaging

#### Multispectral IHC-PhenoImager HT 7-color

Through our collaboration with the Human Immune Monitoring Shared Resource (HIMSR) at the University of Colorado School of Medicine we performed multispectral imaging using the PhenoImager HT instrument (formerly Vectra Polaris, Akoya Biosciences). To quantify levels of microglia, T cells, IL15-GFP, and neurons, formalin-fixed paraffin-embedded tissue sections were stained consecutively with specific primary antibodies for CD45, CD3, CD8, TMEM119, NeuN, and GFP. Briefly, the slides were deparaffinized, heat treated in antigen retrieval buffer, blocked, and incubated with primary antibodies, followed by horseradish peroxidase (HRP)-conjugated secondary antibody polymer, and HRP-reactive OPAL fluorescent reagents. The slides were stripped between each stain with heat treatment in antigen retrieval buffer. Whole slide scans were collected with PhenoImager HT v2.0.0 software using the 20x objective with a 0.5 micron resolution. Spectral references and unstained control images were measured and inForm software v3.0 was used to create a multispectral library reference. The whole slide images (qptiff files) were spectrally unmixed using PhenoImager HT v2.0.0 software (.unmixed.qptiff files). The images were analyzed with tissue segmentation, cell segmentation, and phenotyping using inForm software v3.0 (Akoya Biosciences) and data were compiled and summarized using PhenoptrReports (Akoya Biosciences.

To quantify levels of cleaved caspase 3 and phosphorylated Stat1, formalin-fixed paraffin-embedded tissue sections were stained consecutively with specific primary antibodies for Cleaved Caspase 3, pStat1, GFAP1, Neun, Iba1, SMA, and fibronectin. Briefly, the slides were deparaffinized, heat treated in antigen retrieval buffer, blocked, and incubated with primary antibodies, followed by horseradish peroxidase (HRP)-conjugated secondary antibody polymer, and HRP-reactive OPAL fluorescent reagents. The slides were stripped between each stain with heat treatment in antigen retrieval buffer. Whole slide scans were collected with PhenoImager HT v2.0.0 software using the 20x objective with a 0.5 micron resolution. Regions of interest were selected and rescanned using the 20x objective and the multispectral imaging cube.

Spectral references and unstained control images were measured and inForm software v3.0 was used to create a multispectral library reference. The multispectral images (.im3 files) were spectrally unmixed analyzed with tissue segmentation, cell segmentation, and phenotyping using inForm software v3.0 (Akoya Biosciences) and data were compiled and summarized using PhenoptrReports (Akoya Biosciences.

